# NADES as Biocompatible Media for Thermally Stable RNA Molecules

**DOI:** 10.1101/2025.07.23.665770

**Authors:** Lamya Al Fuhaid, Shahryar Khattak, Arwa Alghuneim, Imed Gallouzi, Young Hae Choi, Robert Verpoorte, Geert-Jan Witkamp, Andreia Farinha

**Affiliations:** Biological and Environmental Sciences and Engineering, King Abdullah University of Science and Technology, Thuwal, Saudi Arabia; Institute Biology Leiden, Leiden University, Leiden, The Netherlands

## Abstract

The inherently low thermal stability of RNA poses operational, accessibility, and financial challenges for RNA research and RNA-based technologies. Natural deep eutectic solvents (NADES) have been shown to enhance the stability of various macromolecules, including DNA and protein. This study explores NADES as alternative storage media for RNA preservation and evaluates their biocompatibility in human cells. *In vitro*-transcribed mRNA was stored in various NADES at temperatures ranging from 4−50 °C, and the integrity was assessed by quantifying its translatability, represented by protein-expressing cells. NADES effectively preserved RNA integrity, retaining over 50% of its translatability for at least four months at room temperature (21 °C) and 48 hours at 50 °C, in comparison to −80 °C storage. Notably, the preservation efficacy remained unaffected by temperature fluctuations. Furthermore, the selected NADES concentrations maintained ∼ 99% cell viability, demonstrating their biocompatibility. These findings establish NADES as efficient, biocompatible RNA storage media, enabling stable storage and transport at ambient and extreme temperatures while withstanding sudden fluctuations. This enhanced stability simplifies and expands the accessibility of RNA-based applications. Additionally, the biocompatibility of NADES supports their potential use in RNA-based biomedical applications.

## 1. Introduction

The stability of RNA is critical for RNA-related research and downstream applications. Despite a few variations, DNA and RNA are composed of similar building blocks and have similar structures. As opposed to the double-stranded structure of DNA, RNA is often single-stranded. The inter-strand hydrogen bonding in DNA can lower the total energy and protect the side chains from reactions^1^. Additionally, RNA contains ribose sugar that has an additional hydroxyl group compared to the deoxyribose sugar in DNA. The hydroxyl group can become deprotonated in basic pH conditions, generating an oxygen nucleophile^2^. The oxygen nucleophile may attack the adjacent phosphorus, breaking the phosphodiester bond in the RNA backbone, resulting in RNA hydrolysis^2^. Because of their low stability, RNA molecules are conventionally stored at low temperatures (approximately −80 °C) to maintain their integrity.

Nowadays, messenger RNA (mRNA) vaccines are being developed for a myriad of infectious diseases. The efficacy of mRNA vaccines on mitigating diseases has been recently demonstrated during the COVID-19 pandemic. The potential of mRNA vaccines goes far beyond infectious diseases as they are currently employed for cancer immunotherapy^3^. They are also being developed for other deadly diseases, including human immunodeficiency virus (HIV) and malaria^4,5^. Despite the robustness of mRNA vaccines, the fragility of RNA molecules requires special freezing units of extremely cold temperatures (−90 to −60 °C).

For example, according to the European Medicine Agency (EMA), two key COVID-19 mRNA vaccines, Moderna and BioNTech, are stable for 6 months in frozen conditions, but only up to 6 h at room temperature (RT)^6^.The need to store vaccines in frozen conditions complicates their transport and administration processes. Additionally, the ultra-cold storage units are expensive, complicated to use, and unequally accessible. As a result, a large percentage of mRNA vaccines lose their functionality during transportation or due to improper storage. These barriers create an imparity in the distribution of vaccines, putting developing countries at risk of not receiving proper health care.

Several methods have been developed to improve the stability of pharmaceutical RNA formulations. For example, it has been shown that the 5’ linkages of uridine residues in the mRNA molecule are particularly susceptible to degradation^7^. Thus, uridine residues in *in vitro*-transcribed mRNA molecules have been substituted with pseudouridine or N_1_-methyl-pseudouridine^7^. Alternatively, uridine can be depleted through guanine/cytosine (G/C) content increase, which can consequently increase the thermodynamic stability of the mRNA^7^. Nevertheless, some chemical modifications can alter the biological properties of the molecule, which may introduce immunogenicity. Additionally, most studies investigating chemically modified mRNA focus on examining their cellular and in vivo performances without evaluating their storage stability or shelf life. Moreover, despite the implementation of pseudouridine substitution in some COVID-19 mRNA vaccines, their stability at RT is still limited.

Deep eutectic solvents (DES; singular and plural) are green solvents formulated from non-toxic, low-cost materials^8,9^. They are composed of hydrogen-bonded components, forming low-melting-point liquids with low moisture content^10^. DES are thermally stable, non-volatile, biodegradable, and easy to prepare^8,9^. They have unique supramolecular structures and properties, linking them to diverse applications. For example, DES have been used in the fields of catalysis^11^, extraction^12–15^, and dissolution of pharmaceutical ingredients^16^. DES made from natural metabolites, like sugars, amino acids, and organic acids, are subtyped as natural DES (NADES)^17^. The components of NADES are mostly accessible in food-grade quality. NADES are often biocompatible and have a lower environmental impact than other DES, presenting a superior option for biomedical and environmental applications^18^.

Due to their wide polarity range, DES can dissolve and stabilize various compounds, such as rutin^13^, paclitaxel (Taxol)^14^, and macromolecules, including DNA^19–21^, protein^15,22,23^, and starch^24^. The stabilization capacity of DES is partially attributed to their hydrogen bond network, which lowers the free energy and prevents macromolecule aggregation. The low water activity in DES was also linked to macromolecule stability^25^. An example of macromolecule stabilization was demonstrated by the capacity of choline chloride-glycerol and choline chloride-ethylene glycol DES to maintain DNA for six months at RT^21^. It was proposed that the cations in choline chloride could have participated in the solvation process by interacting with the negatively charged phosphate groups^10,21^.

In this work, NADES are proposed as an alternative medium for storing RNA at RT. The stability of *in vitro*-transcribed model RNA stored in NADES at temperatures ranging from 4−50 °C is examined. Additionally, the translatability of the mRNA *in cellulo* at multiple time points after being stored in NADES is evaluated. The proposed solution does not include additional chemical modifications to the RNA or active alteration of its secondary structure.

## 2. Methods

### 1.2. NADES preparation and characterization

NADES combining betaine (NADES 1−5) and choline chloride (NADES A−E) with glycerol and/or disaccharides were prepared using the heating method reported by Y. Dai *et al*., 2013^14^. The stability of the resulting samples was evaluated over time and only those that did not exhibit crystallization within a week of formation were selected. All NADES were autoclaved prior to their use in tissue culture experiments.

### 2.2. Cell culture and viability test

Human HeLa cells were grown in Dulbecco’s modified Eagle’s medium (DMEM; 4.5 g/L D-glucose and GlutaMAX, GIBCO) supplemented with 10% (v/v) fetal bovine serum (FBS, Sigma), 100 U/mL penicillin and 100 μg/mL streptomycin (GIBCO), according to standard protocols. Maximum working concentrations of NADES were determined based on a viability test in HeLa cells. Cells were seeded in 24-well plates at a density of 3.0×10^4^ cell/well. When the cells reached ∼ 70% confluence, they were treated with varying concentrations of NADES. Cell viability was assessed 24 h post-treatment using the Invitrogen™ EVOS™ Digital Color Fluorescence Microscope and compared to untreated controls. Cytotoxicity at the selected concentration was further evaluated by quantifying the percentage of viable cells via flow cytometry (Becton Dickinson LSRFortessa™) 24 h post-treatment.

### 2.3. RNA *in vitro* transcription

A plasmid encoding the mCherry open reading frame (ORF), including a T7 promoter, poly-A tail, and T7 terminator (Figure S1), was linearized with SacI-HF (New England Biolabs) overnight and purified using a PCR purification kit (QIAGEN). The linearized plasmid served as a template for *in vitro* transcription using the HighYield T7 ARCA mRNA Synthesis Kit (m5CTP/Ψ-UTP; Jena Biosciences), following the manufacturer’s instructions, to generate 5-methylcytidine and pseudouridine-modified mCherry mRNA. The transcribed RNA was purified using the Monarch RNA Cleanup Kit (New England Biolabs), aliquoted on ice, and stored at −80°C. The undigested plasmid, linearized plasmid, *in vitro*–transcribed mRNA, and polyadenylated mRNA were visualized on a 1.2% agarose gel (Figure S2).

### 2.4. mRNA transfection

HeLa cells were seeded in six-well plates at a density of 1.2×10^5^ cells/well and transfected 48 h later with 250 ng mCherry mRNA using Lipofectamine MessengerMAX™ Transfection Reagent (Thermo Fisher Scientific), following the manufacturer’s instructions. Cells were harvested and fixed 18 h post-transfection. The percentage of mCherry-expressing cells was quantified via flow cytometry (Becton Dickinson LSRFortessa™) using a Texas Red filter..

## 3. Results and discussion

### 3.1. NADES exhibit low cytotoxicity in human cell culture models

HeLa cells were cultured and treated with ten different concentrations of NADES, ranging from 2–20 µg/mL, and observed after 24 h. The maximum concentrations that did not result in a visible reduction in the cell viability were selected. The percentage of viable cells was then quantified by flow cytometry and was above 98% (Figure 1). These results confirm the general biocompatibility of NADES with human cell culture models, in line with previous reports^18^. They also demonstrate the specific compatibility of the tested formulations and concentrations. Collectively, the findings suggest that NADES may be safe for use in biomedical applications.

**Figure 1.**
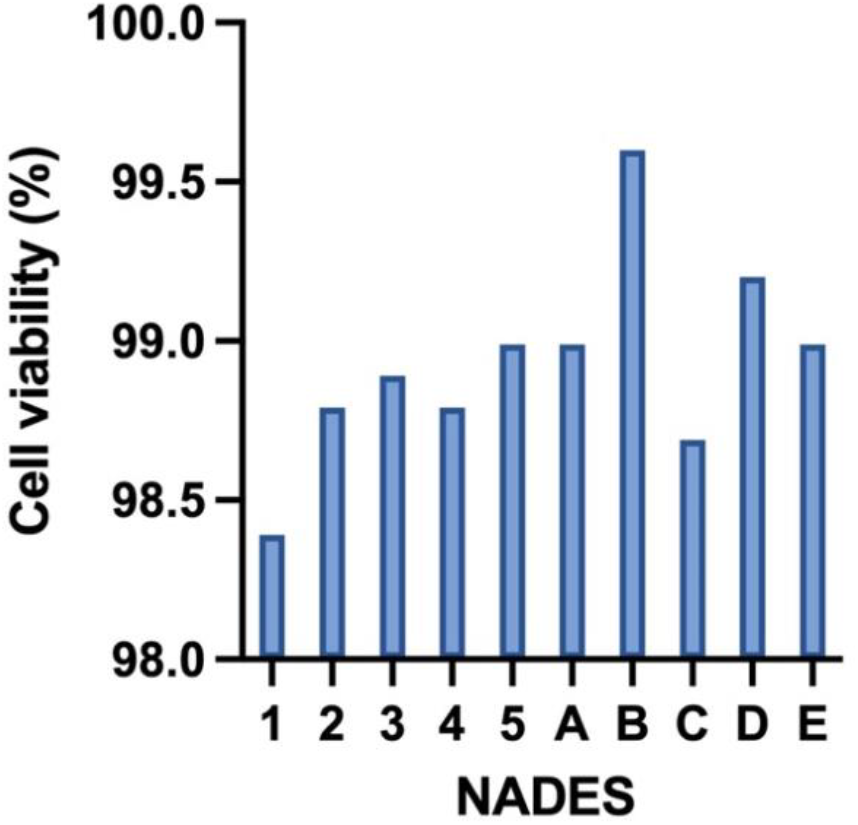
Cell viability HeLa cells treated with NADES. Values are presented as mean ± SD for n=3.

### 3.2. NADES does not interfere with the *in cellulo* translation of mRNA

To assess the ability of NADES to enhance the stability of mRNA without interfering with its translatability, the translation of mRNA into functional proteins after storage in NADES was evaluated. *In vitro*-transcribed mCherry mRNA was used as a model, as it encodes a red fluorescent protein, offering an easy detection and quantification through fluorescence measurement. mCherry is a widely used monomeric fluorophore model protein that absorbs light between 540–590 nm and emits in the 550–650 nm range. The *in vitro*-transcribed mCherry mRNA was stored in water and NADES at 37 °C, a biologically relevant temperature. The *in cellulo* translatability of the mRNA was evaluated after 48 h of storage at 37 °C. HeLa cells were transfected with the mRNA and the percentage of cells expressing mCherry protein was quantified by flow cytometry. Results were normalized to cells transfected with control mRNA stored at −80 °C.

NADES based on betaine (NADES 1−5) were ineffective as mRNA storage media, interfering with the mRNA integrity and/or translatability (Figure 2). Conversely, most of the tested choline chloride-based NADES (A, C, D, and E) supported efficient translation. NADES B was extremely viscous, which complicated handling and may have contributed to the low translation level observed after storage. These results indicate that different NADES formulations have varying effects on mRNA stability, with choline chloride-based systems demonstrating superior retention of integrity and functionality. Because NADES C showed the highest translation level, it was selected for the following experiments.

**Figure 2.**
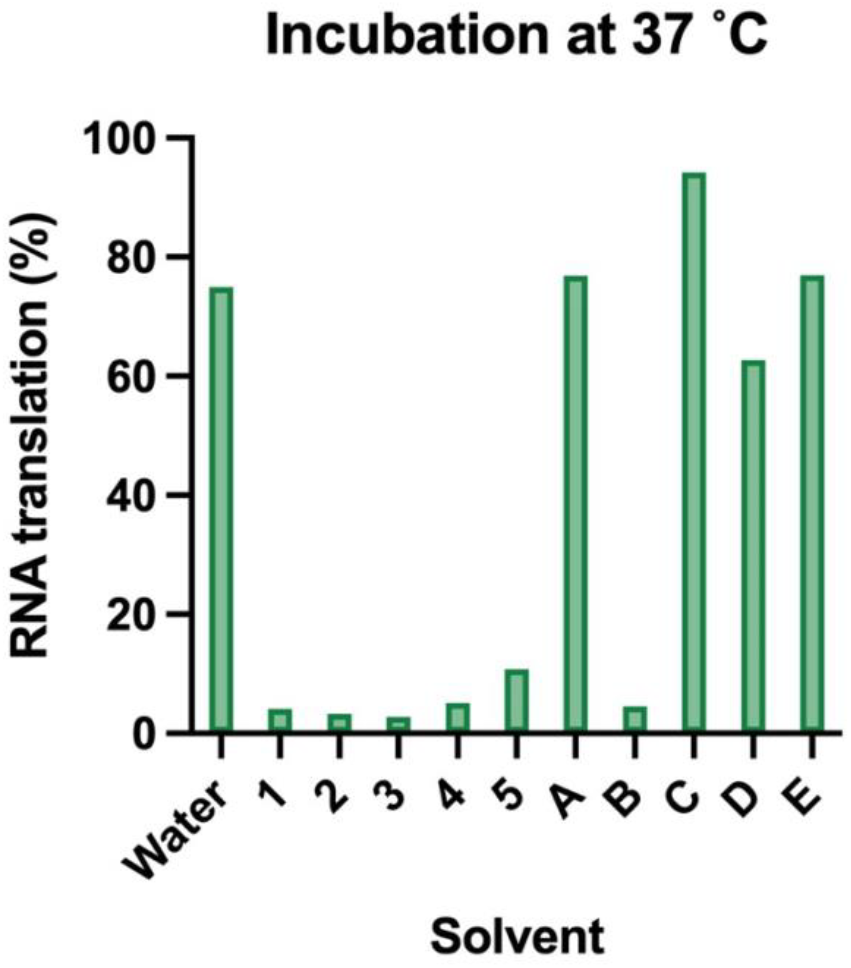
mRNA translation after 48 h incubation at 37°C. mCherry mRNA was incubated in water and NADES 1−5 and A−E, then transfected in HeLa cells. The percentage of cells expressing mCherry protein was quantified by flow cytometry and normalized to the percentage from cells transfected with control mCherry mRNA stored at −80 °C.

### 3.3. NADES protects RNA from degradation at elevated temperatures

The integrity and *in cellulo* translatability of the mRNA was evaluated after 24 and 48 h of storage at 37 °C. HeLa cells were transfected with the mRNA, and the percentage of mCherry-positive cells was normalized to the percentage of positive cells transfected with control mRNA stored at −80 °C. Compared to the control, there was a slight drop in the percentage of positive cells by ∼ 10 and 15% for mRNA stored in NADES and water, respectively after storage for 24 h at 37 °C (Figure 3). An additional 24 h of storage at the same temperature did not cause further reduction. Notably, the used mRNA includes a modified bases, resembling the mRNA molecules currently used in vaccines. This modification infers added stability to the mRNA, reducing its degradation at slightly elevated temperatures.

**Figure 3.**
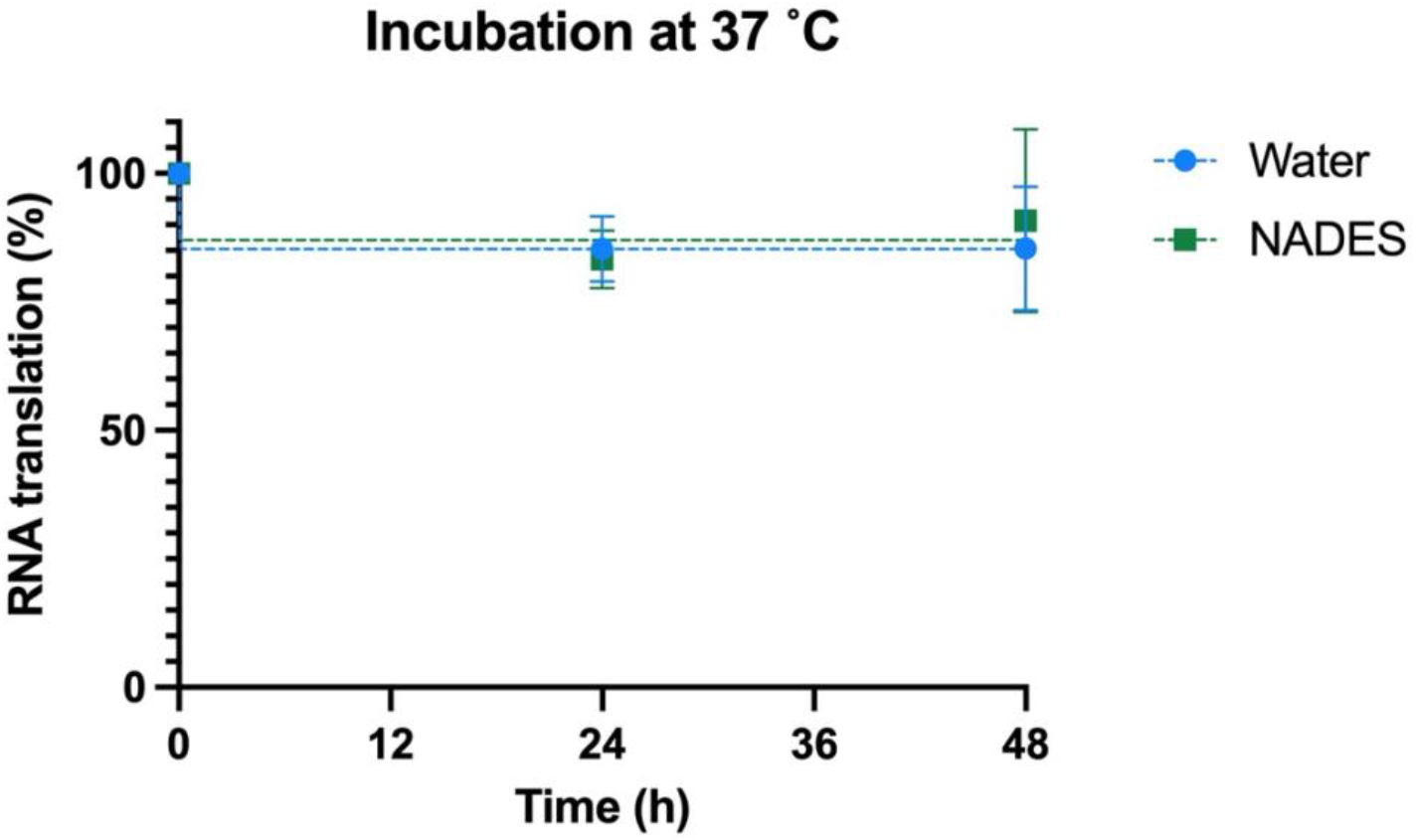
mRNA translation after incubation at 37°C. mCherry mRNA was stored in water and NADES at 37 °C then transfected into HeLa cells. The percentage of fluorescent cells was quantified and normalized to transfection with control mRNA stored at −80 °C. Values are presented as means ± SD for n=3, and the dashed lines represent non-linear one-phase decay fit curves.

Moreover, to test the ability of NADES to protect RNA under extreme conditions, mRNA was also stored at 50°C. This temperature was selected to simulate harsh environments, such as those encountered during vaccine transport, to test the efficiency of NADES as a vaccine transportation medium. When stored in water, mRNA lost more than half its functionality after 24 h and dropped to ∼ 15% after 48 h compared to control (Figure 4). In contrast, mRNA stored in NADES retained ∼ 70 and 50% of its functionality after 24 and 48 h storage at 50 °C, respectively. Collectively, these results highlight the superior protective effect of NADES over conventional storage media and demonstrate their remarkable ability to preserve RNA integrity even under extremely harsh environmental conditions.

**Figure 4.**
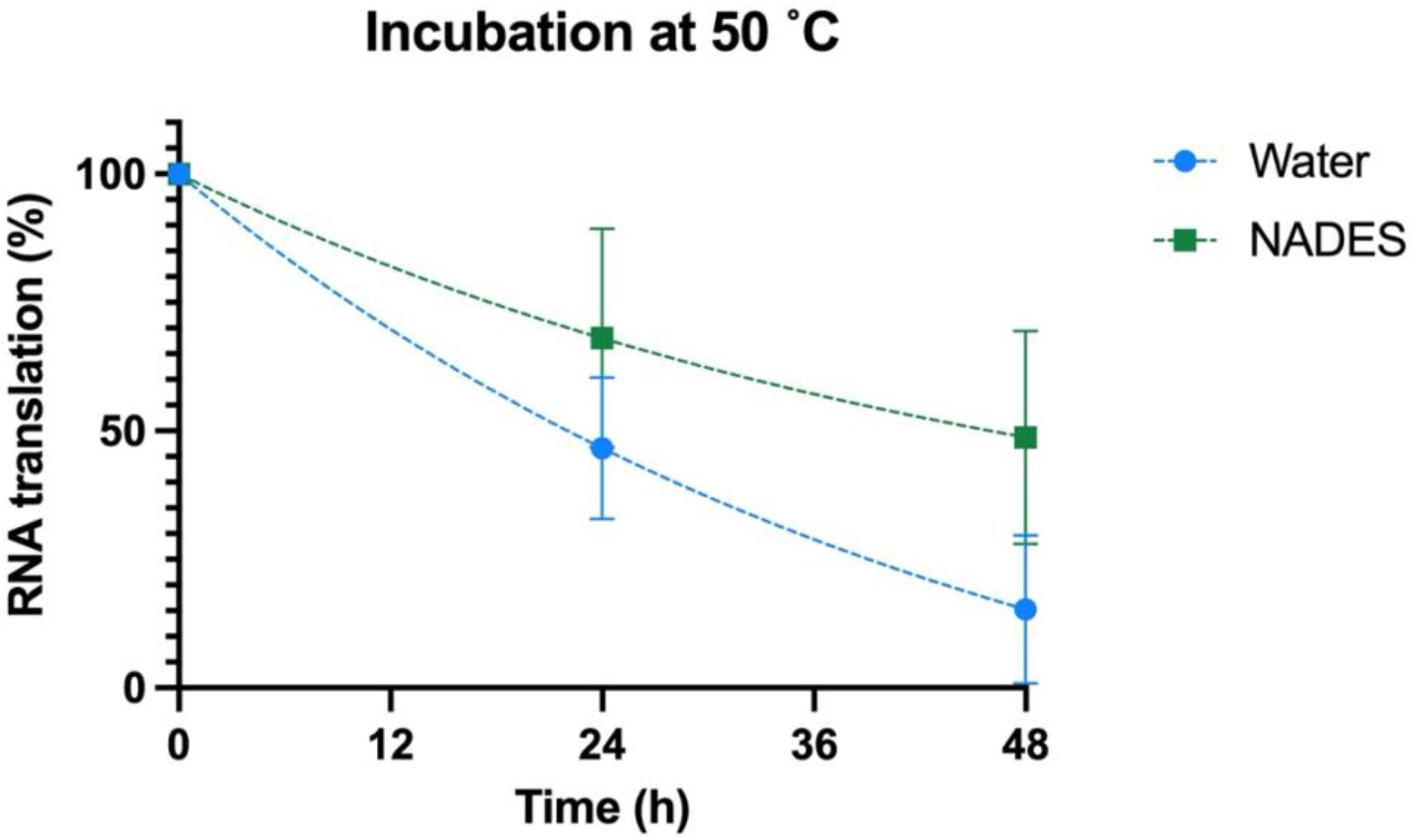
mRNA translation after incubation at 50°C. mCherry mRNA was stored in water and NADES at 50 °C then transfected into HeLa cells. The percentage of fluorescent cells was quantified and normalized to transfection with control mRNA stored at −80 °C. Values are presented as means ± SD for n=3, and the dashed lines represent non-linear one-phase decay fit curves.

### 3.4. Temperature fluctuations do not affect RNA stored in NADES

During a realistic transportation scenario, temperature fluctuations can occur abruptly. To examine the ability of NADES to protect RNA from such changes, mRNA was subjected to four temperature shifts: 12 h at 50 °C, 12 h at 4 °C, followed by another 12 h at 50 °C, and a final 12 h at 4 °C. When RNA was stored in NADES, there was a slight drop in translation after the first 50 °C exposure, but levels remained largely stable afterward (Figure 5). In contrast, RNA stored in water lost almost all functionality after the initial 50 °C exposure. These results clearly demonstrate that drastic temperature fluctuations do not compromise the integrity of mRNA stored in NADES. This underscores the strong potential of NADES for effectively protecting RNA-based formulations and vaccines during transportation, even under harsh and unpredictable conditions.

**Figure 5.**
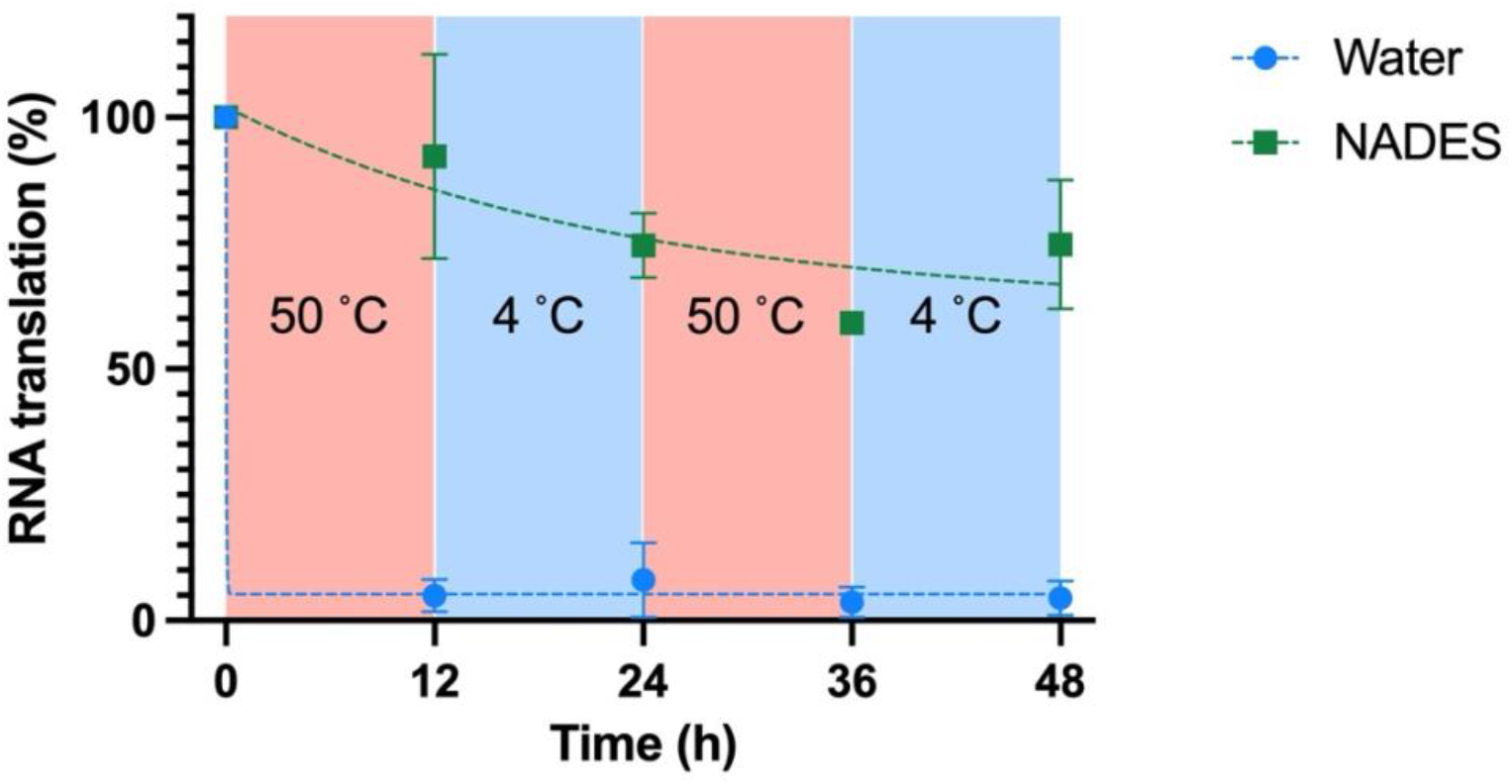
mRNA translation after fluctuating temperature storage. mCherry mRNA was stored at fluctuating temperature for a total of 48 h and evaluated at 12-h intervals. The mRNA was transfected into HeLa cells, and the percentage of fluorescent cells was quantified. Data are normalized to transfection with control mRNA stored at −80 °C. Values are presented as means ± SD for n=3, and the dashed lines represent non-linear one-phase decay fit curves.

### 3.5. mRNA remains stable in NADES at room temperature for extended periods

To evaluate the suitability of NADES as storage media for mRNA molecules and vaccines, the long-term stability of mRNA in NADES was evaluated. The mRNA translation in HeLa cells was evaluated after four months of storage at RT (21 °C). Results show that more than 50% of the mRNA translation was maintained after four months (Figure 6). These results suggest that NADES can eliminate the need for refrigeration during the long-term storage of RNA molecules and RNA-based therapeutics. The ability to preserve such formulations at RT offers significant advantages for research, clinical applications, and public health.

**Figure 6.**
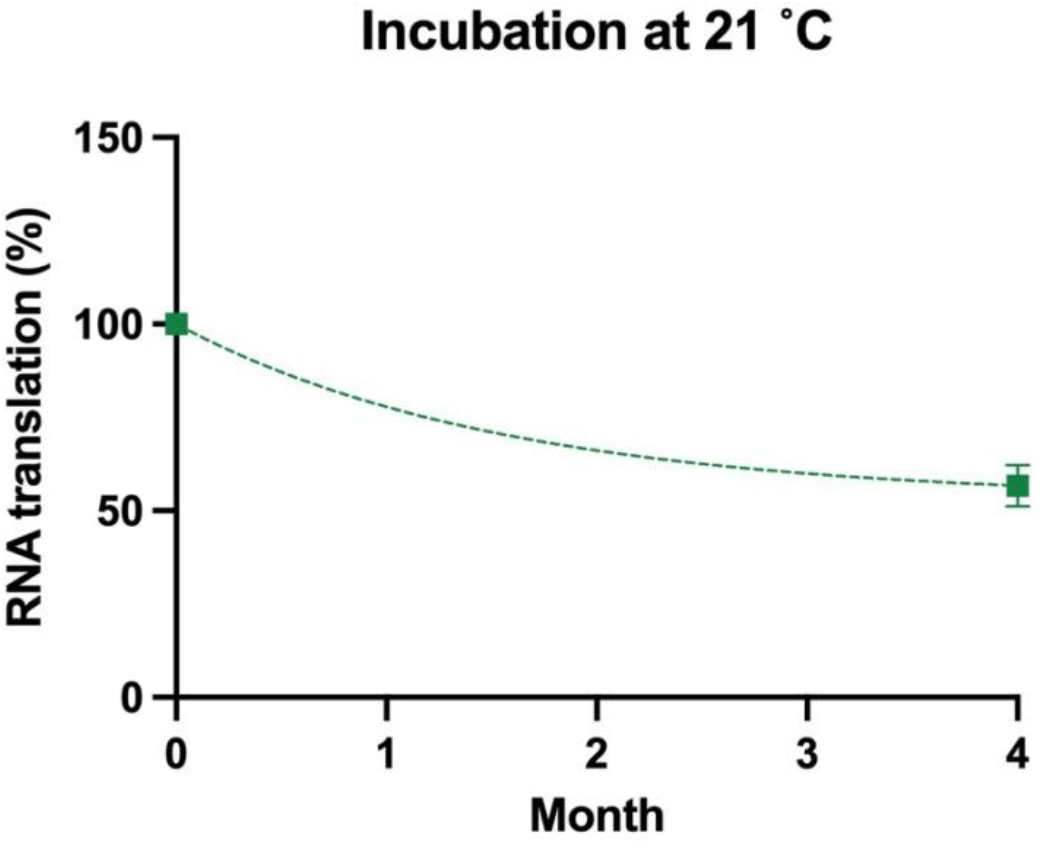
mRNA translation after long-term storage at room temperature (21°C) mCherry mRNA was stored in NADES for four months then transfected into HeLa cells. The percentage of fluorescent cells was quantified and normalized to transfection with control mRNA stored at −80 °C. Values are presented as means ± SD for n=3, and the dashed line represents non-linear one-phase decay fit curve.

## 4. Conclusions

- NADES exhibit low cytotoxicity in human cell models.
- Different NADES have varying effects on mRNA stability and function.
- NADES do not interfere with the mRNA *in cellulo* translation.
- NADES preserved over 50% of RNA functionality *in cellulo* for at least 48 h at 50 °C and 4 months at RT.
- Temperature fluctuations do not affect the stability of RNA stored in NADES.
- The biocompatibility of NADES positions them as strong candidates for biomedical applications, such as mRNA vacccines.
- NADES may extend the shelf life of mRNA vaccines and eliminate the need for refrigeration, substantially reducing storage and equipment costs.
- In vivo testing of mRNA stored in NADES should be carried out to evaluate potential immunogenicity of the mRNA-NADES formulations.

## Acknowledgment

This work was funded by King Abdullah University of Science and Technology. A provisional patent related to this work has been filed (KAUST Ref#: 2025-069-01).

## References

1. Lesnik, E. A. & Freier, S. M. Relative Thermodynamic Stability of DNA, RNA, and DNA:RNA Hybrid Duplexes: Relationship with Base Composition and Structure. Biochemistry 34, 10807–10815 (2002).

2. and, D.-M. Z. & Taira*, K. The Hydrolysis of RNA: From Theoretical Calculations to the Hammerhead Ribozyme-Mediated Cleavage of RNA. Chem Rev 98, 991– 1026 (1998).

3. Barbier, A. J., Jiang, A. Y., Zhang, P., Wooster, R. & Anderson, D. G. The clinical progress of mRNA vaccines and immunotherapies. Nature Biotechnology 2022 40:6 40, 840–854 (2022).

4. Fischer, W. et al./person-group>. HIV-1 and SARS-CoV-2: Patterns in the evolution of two pandemic pathogens. Cell Host Microbe 29, 1093–1110 (2021).

5. Ganley, M. et al./person-group>. mRNA vaccine against malaria tailored for liver-resident memory T cells. Nature Immunology 2023 24:9 24, 1487–1498 (2023).

6. Uddin, M. N. & Roni, M. A. Challenges of Storage and Stability of mRNA-Based COVID-19 Vaccines. Vaccines (Basel) 9, (2021).

7. Oude Blenke, E. et al. The Storage and In-Use Stability of mRNA Vaccines and Therapeutics: Not A Cold Case. J Pharm Sci 112, 386–403 (2023).

8. Zhang, Q., De Oliveira Vigier, K., Royer, S. & Jérôme, F. Deep eutectic solvents: Syntheses, properties and applications. Chemical Society Reviews 41, 7108– 7146 (2012).

9. Liu, Y. et al./person-group>. Natural Deep Eutectic Solvents: Properties, Applications, and Perspectives. (2018) doi:10.1021/acs.jnatprod.7b00945.

10. Smith, E. L., Abbott, A. P. & Ryder, K. S. Deep Eutectic Solvents (DESs) and Their Applications. Chemical Reviews vol. 114 11060–11082 Preprint at 10.1021/cr300162p (2014).

11. Carolin Ruß & Burkhard König. Low melting mixtures in organic synthesis – an alternative to ionic liquids? Green Chemistry 14, 2969–2982 (2012).

12. Morrison, H. G., Sun, C. C. & Neervannan, S. Characterization of thermal behavior of deep eutectic solvents and their potential as drug solubilization vehicles. Int J Pharm 378, 136–139 (2009).

13. Zhao, B. Y. et al./person-group>. Biocompatible Deep Eutectic Solvents Based on Choline Chloride: Characterization and Application to the Extraction of Rutin from Sophora japonica. ACS Sustain Chem Eng 3, 2746–2755 (2015).

14. Dai, Y., van Spronsen, J., Witkamp, G. J., Verpoorte, R. & Choi, Y. H. Natural deep eutectic solvents as new potential media for green technology. Anal Chim Acta 766, 61–68 (2013).

15. Xin, R. et al./person-group>. A functional natural deep eutectic solvent based on trehalose: Structural and physicochemical properties. Food Chem 217, 560–567 (2017).

16. Aroso, I. M. et al./person-group>. Dissolution enhancement of active pharmaceutical ingredients by therapeutic deep eutectic systems. European Journal of Pharmaceutics and Biopharmaceutics 98, 57–66 (2016).

17. Choi, Y. H. et al./person-group>. Are natural deep eutectic solvents the missing link in understanding cellular metabolism and physiology? Plant Physiol 156, 1701–1705 (2011).

18. Hayyan, M. et al./person-group>. Natural deep eutectic solvents: cytotoxic profile. Springerplus (2016) doi:10.1186/s40064-016-2575-9.

19. Zhao, H. DNA stability in ionic liquids and deep eutectic solvents. Journal of Chemical Technology & Biotechnology 90, 19–25 (2015).

20. Mamajanov, I., Engelhart, A. E., Bean, H. D. & Hud, N. V. DNA and RNA in anhydrous media: Duplex, triplex, and G-quadruplex secondary structures in a deep eutectic solvent. Angewandte Chemie - International Edition 49, 6310–6314 (2010).

21. Mondal, D., Sharma, M., Mukesh, C., Gupta, V. & Prasad, K. Improved solubility of DNA in recyclable and reusable bio-based deep eutectic solvents with long-term structural and chemical stability. Chemical Communications 49, 9606–9608 (2013).

22. Knudsen, C. et al./person-group>. Stabilization of dhurrin biosynthetic enzymes from Sorghum bicolor using a natural deep eutectic solvent. Phytochemistry (2020) doi:10.1016/j.phytochem.2019.112214.

23. Lores, H., Romero, V., Costas, I., Bendicho, C. & Lavilla, I. Natural deep eutectic solvents in combination with ultrasonic energy as a green approach for solubilisation of proteins: application to gluten determination by immunoassay. Talanta 162, 453–459 (2017).

24. Zdanowicz, M., Spychaj, T. & Maka, H. Imidazole-based deep eutectic solvents for starch dissolution and plasticization. Carbohydr Polym 140, 416–423 (2016).

25. Schiraldi, A., Fessas, D. & Signorelli, M. Water activity in biological systems - A review. Pol J Food Nutr Sci 62, 5–13 (2012).

