## Supplemental figures for "NADES as Biocompatible Media for Thermally Stable RNA Molecules"

-
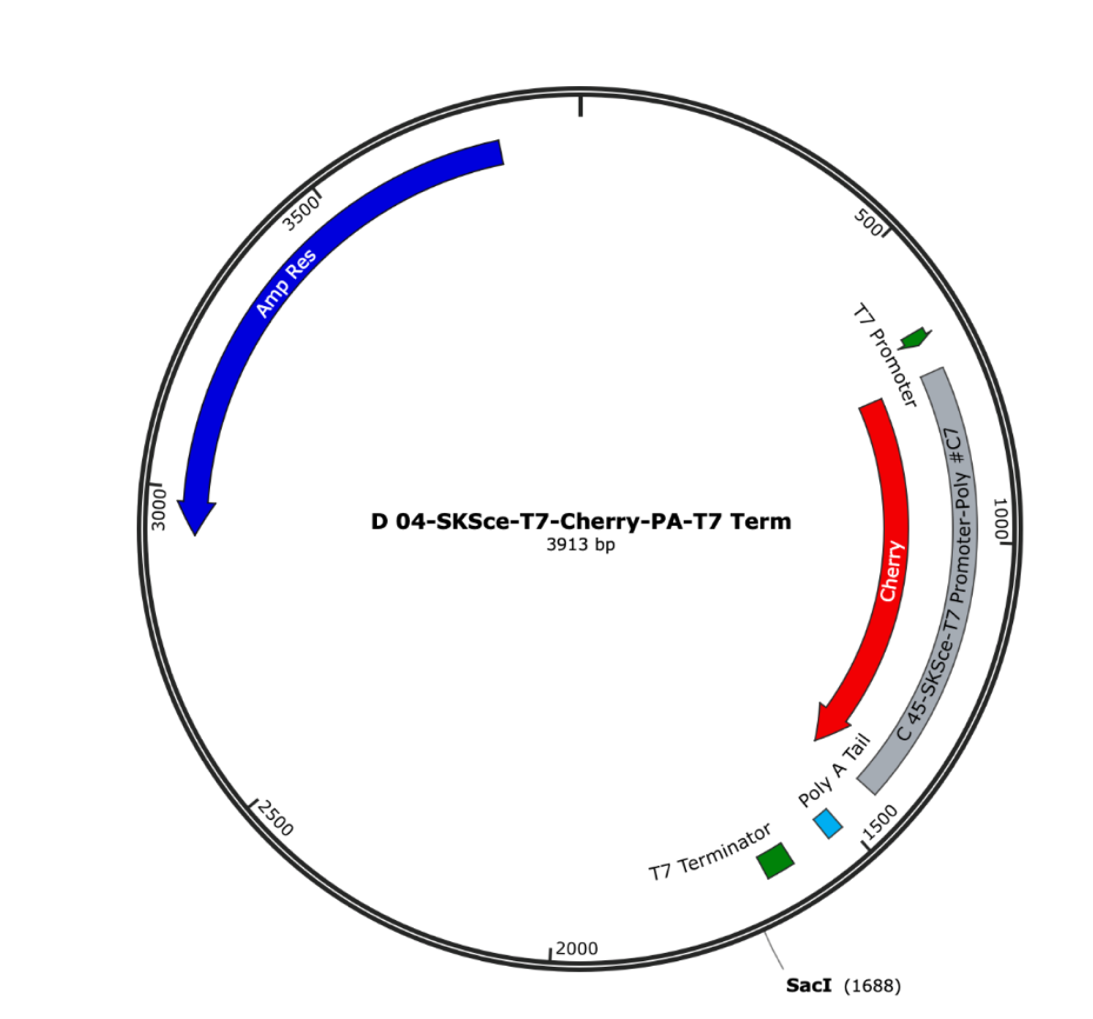

**Figure S1. mCherry plasmid map.** The plasmid containing a T7 promoter followed by mCherry ORF, poly-A tail, and a T7 terminator.

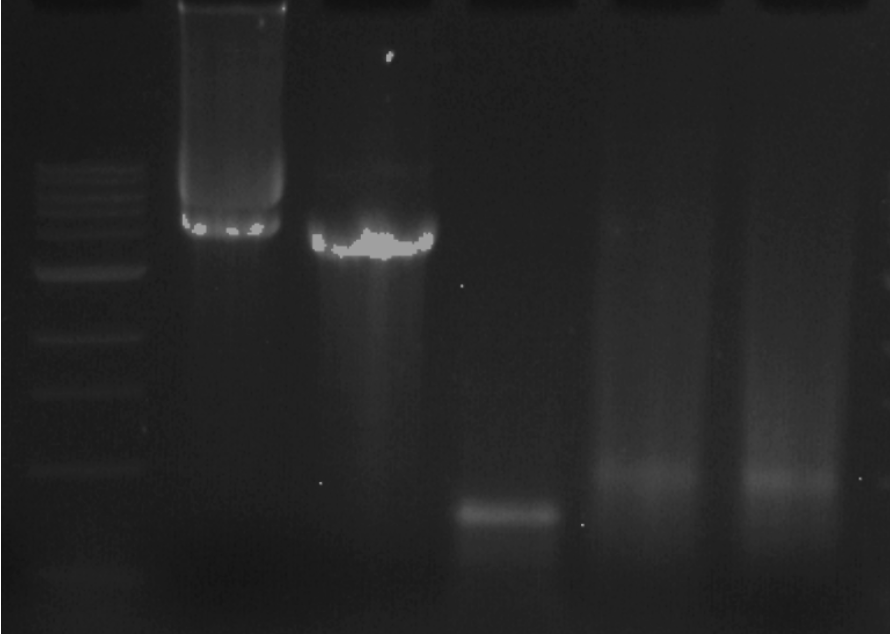

**Figure S2. Gel image conXirming in vitro transcription of mCherry.** From left to right: 1 kb marker, undigested mCherry plasmid, linearized plasmid (digested overnight with SacI enzyme), in-vitro transcribed mCherry mRNA, in-vitro transcribed mCherry mRNA after the addition of a poly-A tail, in-vitro transcribed mCherry mRNA after the addition of a poly-A tail. All samples were run on a 1.2% agarose gel.
